# A Library of Electrophysiological Responses in Plants - A Model of Transversal Education and Open Science

**DOI:** 10.1101/2023.09.29.560074

**Authors:** Danae Madariaga, Derek Arro, Catalina Irarrázaval, Alejandro Soto, Felipe Guerra, Angélica Romero, Fabián Ovalle, Elsa Fedrigolli, Tom DesRosiers, Étienne Serbe-Kamp, Timothy Marzullo

**Affiliations:** Colegio (High School) Alberto Blest Gana, San Ramón, Santiago, Chile; Faculty of Medicine, University of Novi Sad, Novi Sad, Serbia; College of Literature, Science, and Arts, University of Michigan, Ann Arbor, USA; Hirnkastl, Max Planck Institute for Biological Intelligence, LMU Munich, Germany; Backyard Brains, Seoul, South Korea and Ann Arbor, Michigan, USA

## Abstract

Electrophysiology in plants is understudied, and, moreover, an ideal model for student inclusion at all levels of education. Here we report on an investigation in “open science”, whereby scientists worked with students and faculty from Chile, Germany, Serbia, South Korea, and the USA. The students recorded the electrophysiological signals of >15 plants in response to a flame or tactile stimulus applied to the leaves. We observed that approximately 60% of the plants studied showed an electrophysiological response with a delay of ∼3-6 seconds after stimulus presentation. In preliminary conduction velocity experiments, we verified that observed signals are indeed biological in origin, with information transmission speeds of ∼2-9 mm/s. Such easily replicable experiments can serve to include more investigators and students in contributing to our understanding of plant electrophysiology.

## Introduction

All domains of life, from bacteria to humans, use some form of electrical signaling (Johnson et. al 2002, Toyota et. al, 2018, Farmer et. al 2020, Kikuchi et. al, 2022, Adamatzky 2022). Though most commonly associated with the muscle, heart, and nervous system of animals, electrical signals are ubiquitous in life, and, though not as commonly known, also present in plants. The most famous examples of plant electrophysiology deal with rapidly moving plants, such as the Venus Flytrap (*Dionaea muscipula*) (Burdon-Sanderson 1873, Affolter and Olivo 1975, Balotin and Dipalma 1962) and the Sensitive Mimosa (*Mimosa pudica*) (Umrath 1925, Bose 1926, Sibaoka 1962, Hagihara et al. 2022) that generate action potentials when touched. Electrical signaling, however, is also documented in plants that do not necessarily have rapid movement behavior, such as the tomato (*Solanum lycopersicum*) (Reissig et. al 2021), sundew (*Drosera)* (Williams and Pickard 1972) arabidopsis *(Arabidopsis thaliana)* (Mousavi et al. 2013), corn (*Zea mays*) (Kai et al., 2011), avocado (*Persea americana*), and plum (*Prunus domestica*) (RÍos-Rojas et al. 2014), among others.

Plant electrophysiology is understudied and ripe for further experimentation. For example, a PubMed search in 2022 (https://pubmed.ncbi.nlm.nih.gov/) revealed 66 papers published on plant electrophysiology, 1045 on animal electrophysiology, and 2277 on human electrophysiology. Such publication rate difference is curious given that plant electrophysiology experiments are relatively inexpensive and lack regulatory hurdles. Thus, cataloging plant electrophysiology responses is of value to the scientific community, and such experiments are uniquely positioned for the high school and undergraduate teaching environment. Moreover, such improvements in understanding plant electrophysiology could potentially lead to commercially important agricultural yield applications (Reissig et. al 2021). The field is still young, though some companies such as Vivent have begun to integrate such systems into commercial greenhouses.

There is the additional intriguing and provocative hypothesis that if plants have electrical signals similar to properties of nervous systems, therefore perhaps plants are intelligent (Calvo and Lawrence, 2022). Inter-plant communication has been documented in underground root networks (Mahall and Callaway 1991), as well as inter-plant hormone release communication with volatile organic compounds (VOCs) (Shulae et al. 1997, Baldwin et al. 2006). Enabling plant electrophysiological experiments to be accessible at the high school / undergraduate level gives students the ability to contribute to areas of active debate such as learning and information processing in plants (Shin et al. 2021, Gagliano et al. 2016, Taiz et al. 2019), and other concepts such as attention (Parise et al. 2022).

Plant electrical potentials are hypothesized to travel via the plant’s vascular system of xylem and phloem (Nguyen et al. 2018, Rhodes et al. 1996, Wildon et al. 1992, Fromm and Lautner 2007, Choi et al. 2016). Xylem is a structure that transports water and minerals up the plant, while phloem transports bidirectionally, up and down, including the sap from photosynthesis (Lucas et al. 2013). However, non-vascular plants such as moss are also capable of generating electrical potentials (Koselski et al. 2020), which merits further investigation into the anatomical basis of electrical signaling in plants.

With this goal in mind, the work reported here consisted of five high school students (DM, DA, CI, AS, FG), two high school faculty (AR, FO), two undergraduate students (EF, TD), and two scientists (ESK, TM) working together collecting plant electrophysiology data during the 2022-2023 school year. In the specific experiments described in this paper, we recorded from the branches of 16 different plant species in response to flame or tactile stimuli, with 60% of the plants showing electrophysiological responses to the stimuli (examples - **Fig. 1**). Plant electrophysiology is positioned to allow more lateral movement between all levels of education from high schools to advanced research institutes.

**Figure 1-.**
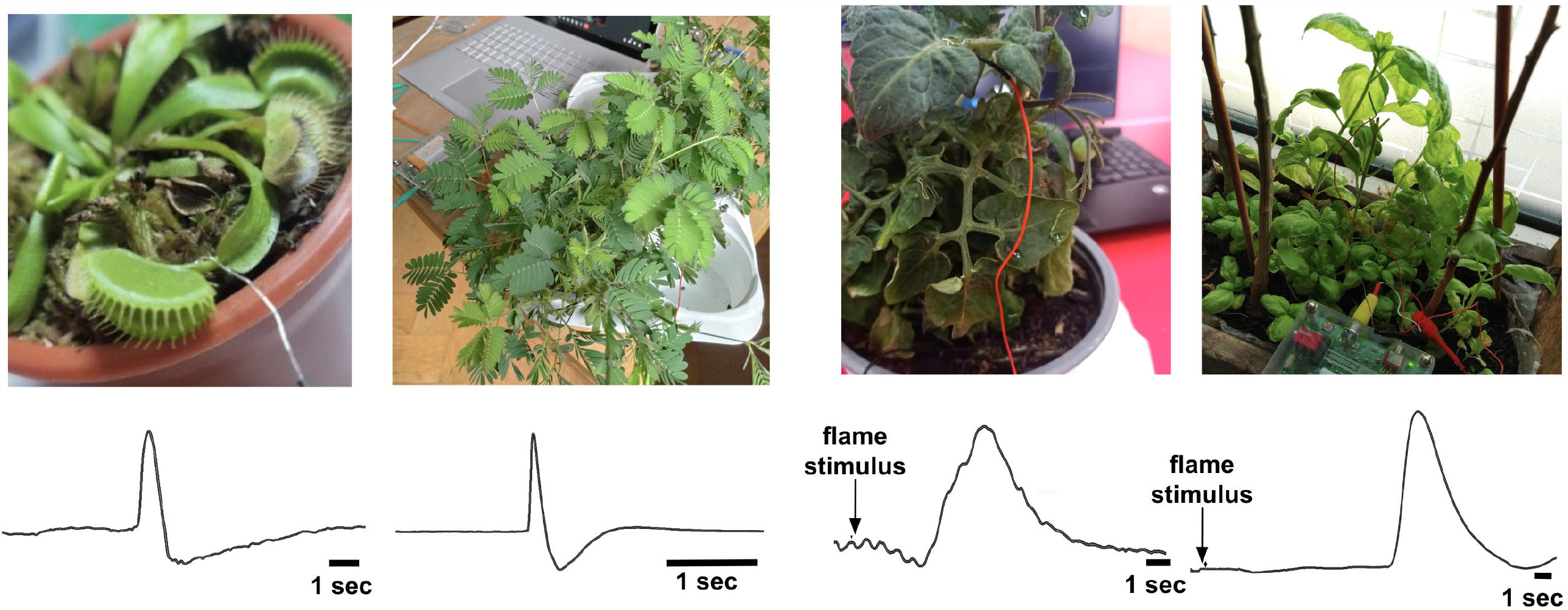
Exemplary Single Trace Dat a from Venus Fly Trap *(Dionaea muscipula)*, Sensitive Mimosa *(Mimosa pudica)*, Tomato *(Solanum lycopersicum)*, and Basil *(Ocimum basilicum)*. The Venus and Mimosa plants received a tactile stimulus, while the Tomato and Basil plants received a flame stimulus.

## Materials and Methods

The electrodes used to record from the plants were 127 μm bare silver wire (A-M systems 781500) wrapped in a spiral 1-3 times around the plant branches, approximately 2-4 cm distal to the leaf being studied. Conductive electrode gel (signa gel, Parker Laboratories, part number #15-60) was applied to the spiral wire to improve signal stability. The ground wire consisted of a standard map pin wire placed into the moist ground of the potted plant (**Fig. 2**). The signals were amplified with a Plant SpikerBox (gain 72x, 0.07-8.8 Hz bandpass filter) (Backyard Brains) and sent via a USB serial interface (10 kHz sampling rate) to a Hewlett Packard, Dell, or Windows Surface laptop computer running on battery power. The software was recorded with the Backyard Brains “SpikeRecorder’’ program as .wav files with event markers stored as .txt files for when the stimulus was applied and removed. Event markers were entered manually by the students during the experiments by pressing the “1” key on the keyboard for when the stimulus was applied and “2” when the stimulus was removed. After each experiment, data was manually uploaded to a Google Drive cloud storage site accessible to all authors.

**Figure 2-.**
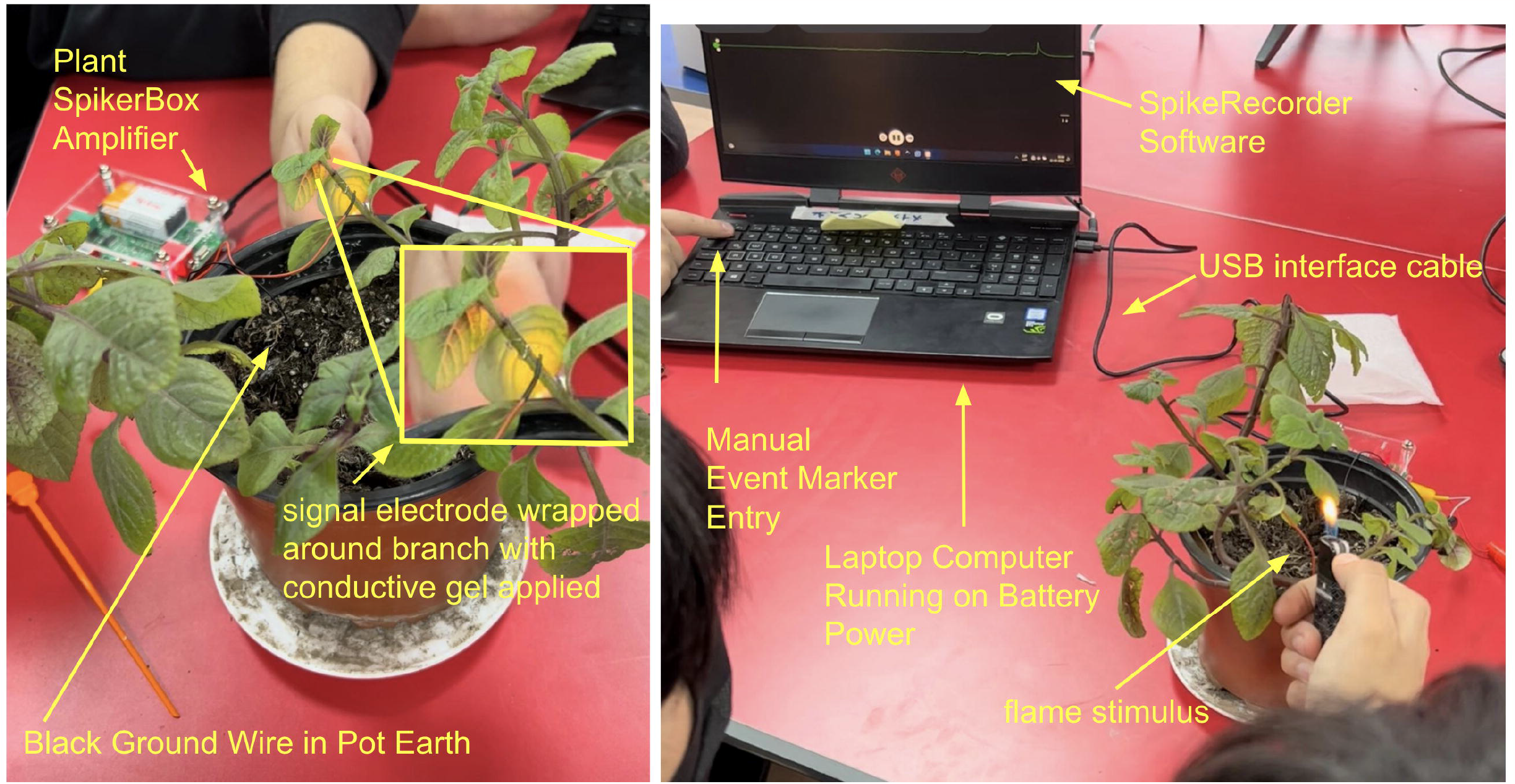
Experimental Setup using Argentinian Dollar *(Plectranthus purpuratus)* as an example. Note student manually marking the point of flame stimulus presentation on the recording software via numerical keystroke.

Experiments were done 80% remotely, with the high school students and faculty (Chile) logging into a Google Hangouts video link, and the two scientists (ESK, Germany, TM, South Korea) guiding the experiments the high school students performed. The last month of the class in Chile was done presentially, with TM visiting the high school for the final classes and ESK logging in remotely. The experiments the undergraduates (EF, TD) did were done during a four-week summer fellowship in Belgrade, Serbia with the scientists ESK and TCM physically present. Effective use of online tools (Google Hangouts, Google Colab, and Google Drive) was crucial for the success of this project.

The plants examined in detail (more than 3 experiments over different days) were 1-Araucaria (*Araucaria araucana*), 2-Argentinian Dollar (*Plectranthus Purpuratus*), 3-Basil (*Ocimum basilicum*), 4-an unidentified fern species (Polypodiopsida), 5-Ivy (*Hedera*), 6-Lemon Balm *(Melissa officinalis*), 7-Mint (*Mentha spicata*), 8-Oregano (*Origanum vulgare*), 9-Papyrus (*Cyperus papyrus*), 10-Radiators (*Peperomia*), 11-Rosemary (*Salvia rosmarinus*), 12 -Ruda (*Ruta graveolens*), 13-Sensitive Mimosa (*Mimosa pudica*), 14-Sundew (*Drosera capensis*), 15-Tomato (*Solanum lycopersicum*), and 16-Venus Fly Trap (*Dionaea muscipula*), all purchased from various local suppliers (**Suppl. Fig. 1, Table 1**). For the Venus Fly Traps, we used tactile stimulation of the trap trigger hairs with a plastic probe as the stimulus. For the Sensitive Mimosa, we used a tactile stimulus of the leaves with a plastic probe. For all other plants we used flame applied to the apical tip of a selected leaf closest to the recording electrode for 2-4 seconds as the stimulus. The flame stimulus consisted of flexible long-neck butane lighters (Ronson) carefully positioned to minimize any movement of the plant leaves and branches during experiments to reduce electrical artifacts. The flame stimulus was chosen as a stimulus to minimize any electrical noise caused by movement of the leaves or contact with metal scissors (cutting stimulus). Typical appearance of the leaf after burning was a charring around the leaf apical tip. All plants studied had a minimum of 2 members of the species studied (such as Fern and Papyrus), while others had up to 9 (such as Mint and Sundew).

In some instances, the electrode technique varied. For the Venus Fly Traps, the recording electrode consisted of a plastic stake placed into the ground with a spiral wrap of silver wire around the tip, and the spiral tip of silver lain against the side of the trap, with conductive gel applied to the silver spiral. For some of the Araucaria, Radiator, and Papyrus experiments, the ground electrode consisted of a stainless steel map pin needle placed directly into the stem instead of the moist soil. Experiments were performed approximately once a week from late August 2022 to March 2023 during the approximate spring and summer seasons in Santiago, Chile, and during a four-week 2023 summer fellowship with undergraduate students in Belgrade, Serbia.

To analyze the data, we used custom Google Colab notebooks that could access our recordings shared in a Google Drive. The software routines were written in Python, and code base examples can be seen in the “Online Resources” links at the bottom of this manuscript. Our “spikertools” library analyzed the plant electrophysiological responses before and after the flame/tactile stimulus was applied using the embedded event markers. All recordings were normalized for signal amplitude, peak-aligned, and sign inverted when the peak response was negative. To determine general response patterns (see **Suppl. Fig. 1**) we calculated the mean over all individual recordings where we detected at least 10 peaks after artifact rejection (**Fig. 3** - the green line represents the mean with standard deviation shades and the gray traces depict individual recordings). Plants were considered to have responses to the stimuli if the recorded signal deviated more than 75% outside the signal range prior to the stimuli. Automated data processing enabled the selection of the data for anticipated signals. We filtered out traces that 1) had spikes faster than 500ms (touching artifacts of the recording wire) 2) were exceeding the recording range (clipping artifacts) 3) were exhibiting spikes that occurred over an elevated baseline level of 75% (normalization artifacts). This analysis allows us to add more recordings to our dataset, but also to extend the automated signal detection for new plant species.

**Figure 3-.**
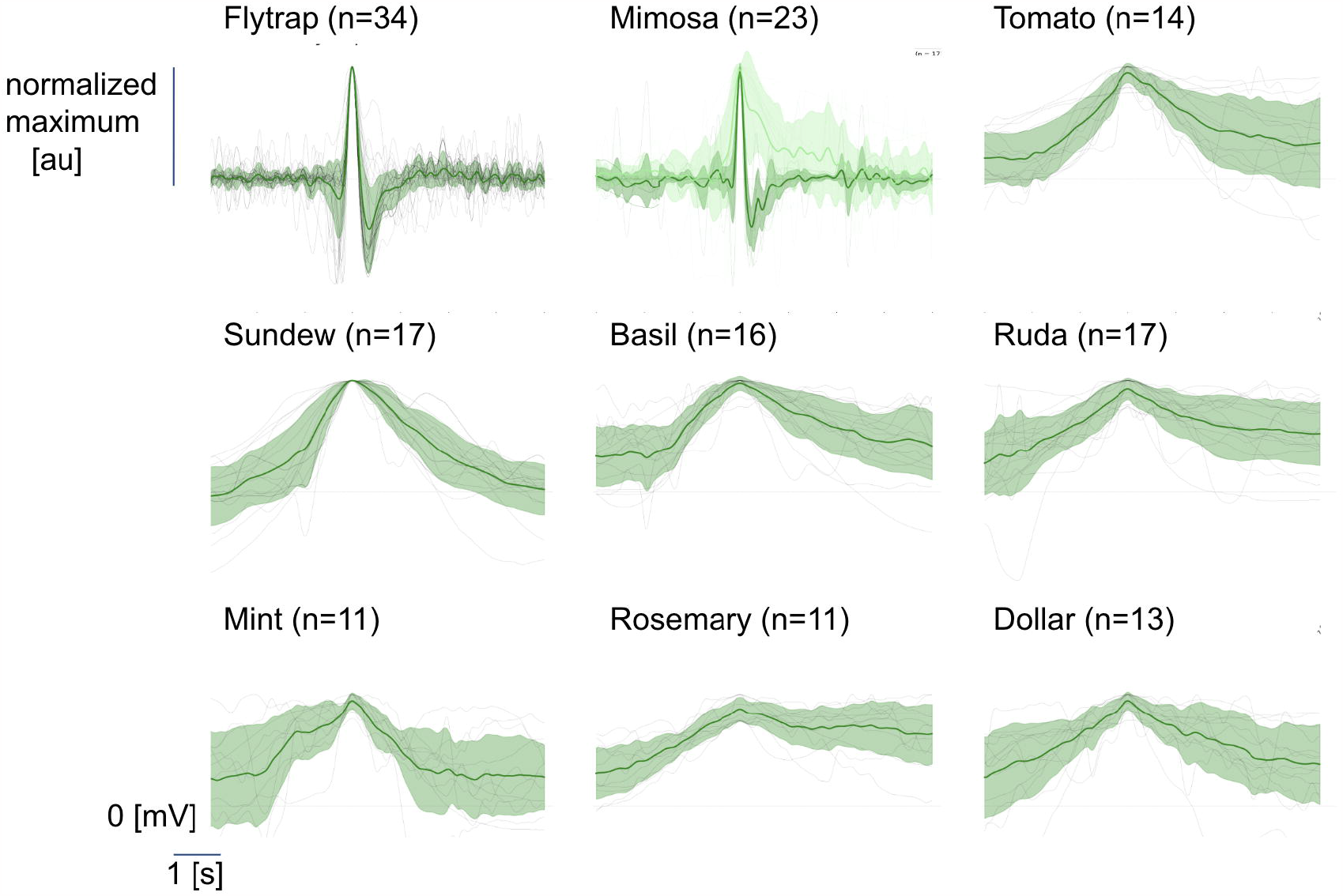
Library of electrical discharges from 9 plants. Robust Flytrap (n=34) and Mimosa (n=31) action potentials elicited by tactile stimuli. Mimosa recordings showed fast (n= 14, green) and slow (n=9, light green) signals. Other plant recordings show putative wound potentials obtained through a flame stimulus. Depending on the stimulus, different data analysis was performed (see methods). Individual grey traces depict single experiments and green lines show the overall mean with the standard deviation as shaded areas. Horizontal dashed lines at O for reference after baseline subtraction. Vertical bar depicts the maximum of the normalized responses.

In some final experiments, we measured the conduction velocities of electrical impulse transmission in the tomato (*Solanum*), sundew (*Drosera*), and sensitive mimosa (*Mimosa*) plants (**Fig. 4**). To record the conduction velocities, we used a 2-channel prototype amplifier that consisted of two custom amplifier shields (bandpass 0.2 - 130 Hz, gain ∼55x) placed on top of an Arduino Uno for signal acquisition. Two wires (channel 1, channel 2) were placed with conductive gel approximately 2-4 cm apart on a branch and a ground pin placed in the ground (sundew / mimosa) or in the stem (tomato) For the tomato and sundew measurements, a flame stimulus was used. For the sensitive mimosas, a tactile stimulus was used (a strike on the leaves with a plastic probe). The distance between the two signal wires on the branch and the time difference between the two electrophysiological peaks on the two channels was used to calculate conduction velocity.

**Figure 4-.**
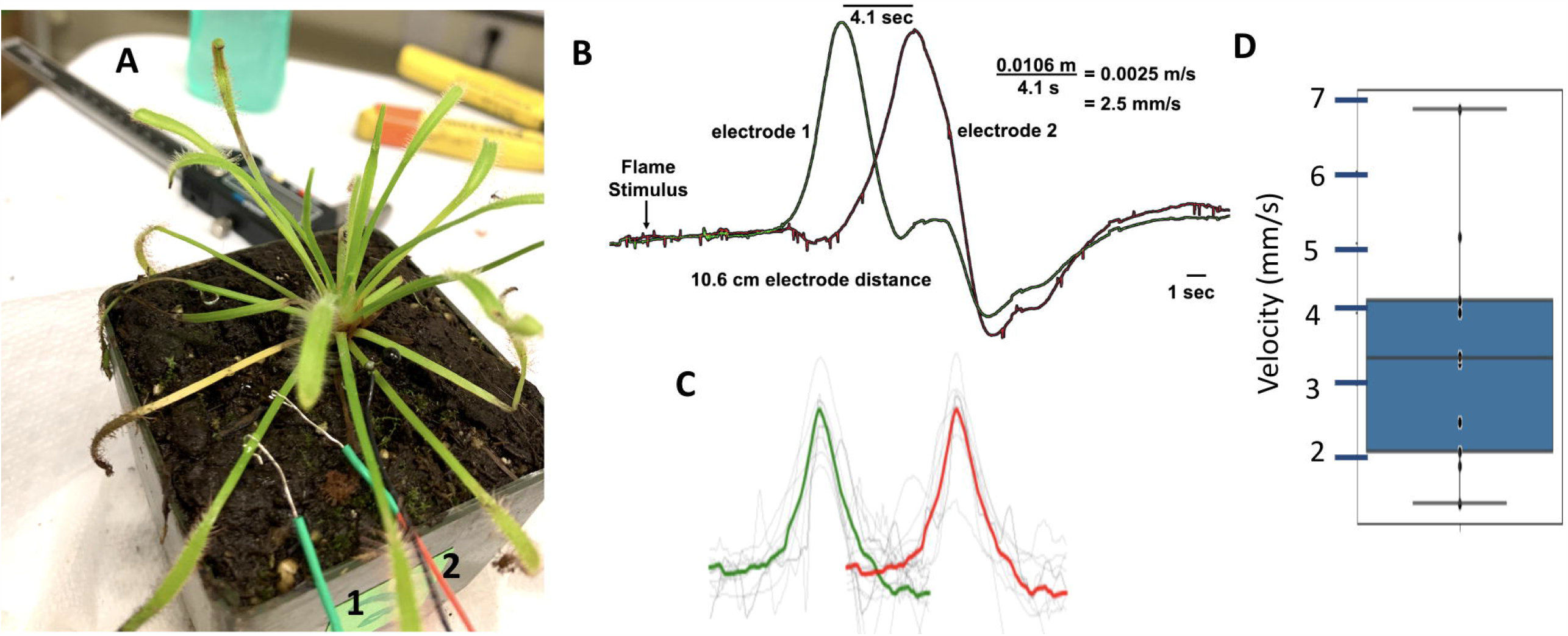
Conduction Velocity Measurements in Sundew *(Drosera capensis)* using a flame stimulus. **A**. Experimental Setup with two signal electrodes along one branch. **B**. Exemplary recording. C. Data and Average Signal from 11 different recordings on 9 plants from two channels (green and red). **D**. Scatterplot of same 11 recordings. Average conduction velocity is 3.2 mm/s.

## Results

**Fig. 1** shows exemplary data (individual traces) from a Venus Fly Trap, sensitive mimosa, tomato and basil plant. The Venus Fly Trap and mimosa show responses to tactile stimuli, while the tomato and basil recordings show responses to flame stimuli. Note the obvious change in signal when the stimulus was applied.

**Fig. 2** depicts the standard experimental set-up used by the students, which they learned rapidly to operate (within one session). Unless otherwise noted in the methods above, a silver wire signal electrode was wrapped around the branch, conductive gel was applied, and a map pin ground wire was placed in the moist earth of the potted plant. The signal was amplified with our custom Backyard Brains “Plant SpikerBox” amplifier, and the electrophysiological responses were recorded using our “Spike Recorder” software.

**Table 1** shows the list of all 16 plants studied and the total normal of recordings obtained. The total number of recordings obtained was 398, taken over a period of on average one session per week during 7 months spread out over one school year (spring-summer - Chile; summer - Serbia).

**Fig. 3** demonstrates the data from the 9 of 16 (56%) plants that demonstrated a putative response to the flame or tactile stimulus. The plants that showed responses include *Ocium* (Basil), *Dionaea* (Venus Fly Trap), *Ruta* (Ruda), *Drosera* (Sundew), *Solanum* (Tomato), *Mimosa* (Mimosa), *Mentha* (Mint), *Salvia* (Rosemary), and *Plectranthus* (Argentinian Dollar). If the amplitude exceeded 75% of the average signal prior to the stimulus, the recording was considered a putative response. Of note is that the responses can be seen in plants (7 of 9) that do not necessarily exhibit rapid movement in their behaviors as well as in common household plants that are available for purchase globally, such as the *Solanum* (tomato), *Ocimum* (basil), and *Mentha* (mint).

**Fig. 4** shows experimental results on conduction velocity measurements in *Drosera* (sundew) in response to a flame stimulus. Conduction velocities were on the order of 2-9 mm/sec. This is of note, as sensory nerve neural conduction velocities in humans is ∼50-80 m/s (Galdames and Castillo 2004) and in the earthworm, 10-20 m/s (Shannon et. al 2014). Thus, plants are transmitting information throughout their vascular structures on an order of 1000-15,000x slower than animals’ neural systems. Preliminary conduction velocity experiments were also done in tomato plants, with an observed conduction velocity of ∼9 mm/s in response to a flame stimulus, and in the sensitive mimosa a conduction velocity of ∼7.7 mm/s in response to tactile stimuli (data not shown).

## Discussion

These experiments enabled students and faculty to see plants as much more dynamic systems than often thought - a theme called “plant blindness” (Wandersee and Schussler 1999). Such experiments are open-ended and readily reproducible, facilitating result verification by scientists and students globally. Moreover, our online data repository welcomes additional observations, fostering collaborative contributions to the field.

In this work we demonstrated and verified that many plants (not necessarily rapid-movement plants) have robust electrical signals. Studying 16 different species is a high number from an individual experiment point of view, but not considering the high biodiversity of plants in the world. The number of plant species is estimated to be 400,000 (Antonelli A et al. 2020), which seems oddly small and potentially underreported, as there are more documented animal species, mostly due to insects (> 900,000 insect species) (Sabrosky 1952). Our hope is that electrophysiology in many diverse plant species becomes commonplace in educational institutions globally, increasing our understanding of information transmission in plants, a subject we have anecdotally found that teachers and students find fascinating.

Though not systematically studied, we also examine other plant species in one-off experiments, such as clover (*Trifolium*), chile (*Capsicum*), touch-me-not (*Impatiens*), rose (*Rosa)*, and carrot (*Daucus)*. We observed signals while stimulating through cutting or flaming different parts of the plants (leaves, fruits, stem). The chile, rose, and touch-me-not signals were compelling, but require more in-depth investigation. Studying the electrophysiological systems of plants valuable for the global food supply (such as tomato, chile, and carrot) has obvious social implications - could electrophysiological detection systems for plants help in greenhouse agriculture for automated health detection (Reissig et. al 2021)?

Plant electrophysiology is paradoxically both easier and harder than animal electrophysiology. It is easier as the experiments are relatively simple to set up, but without a behavior observable on a human time scale, interpreting whether observed electrophysiological data is biologically relevant and real remains the primary challenge. Disregarding noise sources systematically is the main experimental difficulty for novices. By building easily understandable artifact rejection routines into our analysis script, we can reduce bias in data visualization of our recordings that could potentially be caused through “hand sorting” (see **Suppl. Fig. 1** for a representation of all data recorded without artifact rejection applied).

In all plants aside from the Venus Fly Trap and Sensitive Mimosa, the responses were very variable. For newly investigated plants that exhibited potential wound responses, we only detected them in ∼30% of all total recordings. The variability is most likely linked to the health and physiological state of the individual plants used. For example, in mint plants, sometimes, at the same time of day, in the same environmental conditions, one plant would show robust responses to the flame stimuli, while the next plant investigated of the same species would not reveal any responses. Thus we make the statement “putative” to state that though the analysis automatically went through our algorithm, determining the origin of such variability is obviously of interest for future experiments. We found the non-rapid movement plants with the most “robust” and “reliable” wound responses were the basil and tomato plants (**Fig. 1**), a pleasant surprise given how easy it is to purchase these plants in all five countries involved in this study. Interestingly, to our knowledge, we were the first ones to describe the electric potentials of wound responses in the basil plant, whereas secondary effects of wounds are well studied describing phenolic compound production, VOC release, and hormonal responses (Vrakas K et al. 2021).

As all of our data is on an online repository (see “Online Resources” at the bottom of this manuscript), whereby other investigators (professional scientists, students, and citizen-scientists), can add their own experimental data, we hope to develop a large “citizen science” library of plant electrophysiological response data that scientists around the world can access and analyze. It is a non-trivial part of this work that high school students did the majority of the data collection, showing the advantages that plant electrophysiology has for introducing young students to the scientific process of data collection, data analysis, and manuscript writing, something most aspiring scientists are only exposed to in late undergraduate education at the earliest, if lucky. The participation of professional scientists on a consistent, long-term basis with young researchers is critical for such initiatives. Young students can record data and discuss results, but analyzing the literature and the technical writing is the biggest challenge, which requires active oversight and collaboration by scientist colleagues.

We recorded from principally angiosperms (flowering plants); the only gymnosperms were young Araucaria trees, and the only Polypodiopsida was one unidentified fern species. We did not try to record from non-vascular plants such as mosses, though that would be an interesting future direction.

As we observed 56% of the plants we studied had electrical responses to our stimuli, does this mean that the other 44% did not have electrical responses, or merely that we failed to observe them? The latter seems more likely given how ubiquitous electrical signaling is in biology. Effects on the responsiveness of the plant may include the dormancy state of the plants, health of the plant -whether the plant has already been attacked by insects (we anecdotally observed plants with herbivore wounds did not manifest electrophysiology wound responses, but did not systematically study this), moisture level of the soil (ensuring a stable ground), among other factors. Another potential source of noise can be the signal interface, as most of our plants with observed responses were relatively small with slender, pliable green stems (<1 meter tall), whereas other plants such as the papyrus and rosemary had drier stems and branches.

With large plants like trees, would a simple spiral electrode of silver covered with conductive gel be sufficient for recording a signal? For example, in the Araucaria plant, the leaves and branches are very tough and waxy. Was a lack of observed response due to an improper interface with the plant, or that the plant lacks electrical signals? Testing with various different electrode designs would bear this out. Given how ubiquitous electrical signals are across all domains of life, such a lack of electrical response seems unlikely in the plants where we didn’t observe a response. It is more likely due to the failure of improper interface design.

The questions that emerge from this work are: 1) How are these electrophysiological signals generated in the plant?, and 2) How far do these electrical signals propagate? Future experiments will involve systematically measuring conduction velocity across the plants studied in this work to visualize the propagation paths and anatomical structures (**Fig. 4**). An additional question is: what is the function of such signals (Simons 1992)? Plants face the “stuck in place” problem, thus they have developed defenses against attack, most notably in the willow tree, tomato, tobacco, and sensitive mimosa (Chamovitz 2017).

It is our hope that in the future more and more scientists contribute to a database of plant electrophysiology, and we improve our enlightenment of one of the least appreciated, though no less marvelous, domains of life. Our work here, with high school students collecting the majority of the data, speaks to the universal possibility of increased public participation from all realms in the scientific process of understanding biological information-processing systems.

### Online Resources

To generate the figures **Fig. 3** and **Suppl. Fig. 1**, we used the following code base: t.ly/dJBGc

To generate **Fig. 4** (conduction velocity in drosera), we used the following code base: t.ly/HRdQA

The online data from all recordings is stored in the following library: t.ly/imDXR

New data can be sent also through Google Forms: t.ly/LbTTI

All data recorded in this study are available at the permalink: t.ly/BaZhY

## Acknowledgments

Thanks go to “el Don” Ricardo Román, director of high school Colegio Alberto Blest Gana, for enthusiastically blessing this unique educational experience at his institution. Nour Chahin designed initial prototypes of the Python toolbox ‘spikertools’ that we used for our data loading and processing steps. This work was principally supported by subsidies awarded by the government of the Republic of Chile designated for use by public high schools. Additional support was provided by the Center for the Promotion of Science in Serbia and the Nordeus Foundation. We also thank Daniela Flores, Kadeem Gilbert, and Christopher Harris for carefully reading and commenting on our manuscript.

## Conflict of Interest Statement

Authors TD, ESK, and TM work at the company Backyard Brains, which sells the amplifier equipment mentioned in this manuscript. Authors DM and DA were also interns for three months at Backyard Brains during the preparation of this manuscript after the formal class ended.

## Notes

https://t.ly/BaZhY

