## Supplementary figures and images for "A Library of Electrophysiological Responses in Plants - A Model of Transversal Education and Open Science"

### Supplemental Figure 1

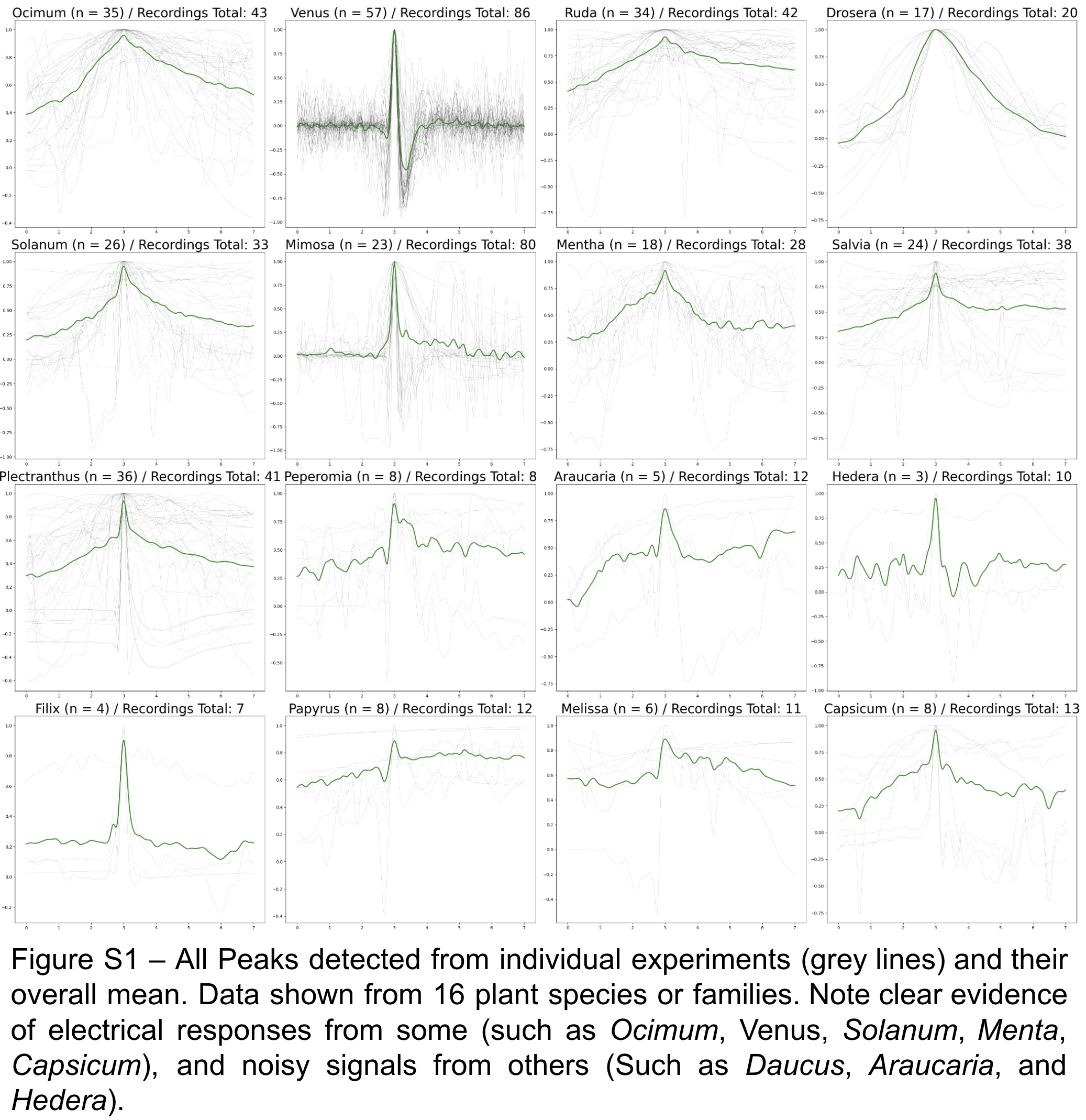
